# Platelet Thrombus Formation in eHUS is Prevented by Anti-MBL2

**DOI:** 10.1101/707364

**Authors:** R. I. Kushak, D.C Boyle, I.A. Rosales, J.R. Ingelfinger, G.L Stahl, M Ozaki, R.B. Colvin, E.F. Grabowski

## Abstract

Epidemic Hemolytic Uremic Syndrome (eHUS) caused by Shiga toxin-producing bacteria is characterized by thrombocytopenia, microangiopathic hemolytic anemia, and acute kidney injury that cause acute renal failure in up to 65% of affected patients. We hypothesized that the mannose-binding lectin (MBL) pathway of complement activation plays an important role in human eHUS, as we previously demonstrated that injection of Shiga Toxin-2 (Stx-2) led to fibrin deposition in mouse glomeruli that was blocked by co-injection of the anti-MBL-2 antibody 3F8. However, the markers of platelet thrombosis in affected mouse glomeruli were not delineated. To investigate the effect of 3F8 on markers of platelet thrombosis, we used kidney sections from our mouse model (MBL-2+/+ Mbl-A/C*−/−;* MBL2 KI mouse). Mice in the control group received PBS, while mice in a second group received Stx-2, and those in a third group received 3F8 and Stx-2. Using double immunofluorescence (IF) followed by digital image analysis, kidney sections were stained for fibrin(ogen) and CD41 (marker for platelets), von-Willebrand factor (marker for endothelial cells and platelets), and podocin (marker for podocytes). Electron microscopy (EM) was performed on ultrathin sections from mice and human with HUS. Injection of Stx-2 resulted in an increase of both fibrin and platelets in glomeruli, while administration of 3F8 with Stx-2 reduced both platelet and fibrin to control levels. EM studies confirmed that CD41-positive objects observed by IF were platelets. The increases in platelet number and fibrin levels by injection of Stx-2 are consistent with the generation of platelet-fibrin thrombi that were prevented by 3F8.

## Introduction

Epidemic hemolytic uremic syndrome (eHUS), the most common cause of acute renal failure in children worldwide, is characterized by the triad of thrombocytopenia, microangiopathic hemolytic anemia, and acute kidney injury, preceded generally by bloody diarrhea. [1, 2]. Shiga toxin (Stx)-producing *E. coli,* especially strain O157:H7, causes the disease; outbreaks of eHUS most often follow infection after ingestion of water, beef products and vegetables contaminated by feces from cattle and other farm animals that carry pathogenic E. coli [3–5] producing one or two major forms of Stx: Stx1 and Stx2. Stx2 is more often associated with disease and is 400-fold more toxic in mice than Stx1 [6]. Acute kidney failure occurs in up to 65% of those who develop eHUS, and many require acute dialysis. Serious involvement of other organ systems may induce gastrointestinal injury and necrosis; pancreatic injury that may lead to diabetes; pulmonary injury and brain injury (neurocognitive disorders, occipital lobe blindness). Death occurs in 3-5% of those affected, and long-term sequelae, including chronic kidney disease and hypertension, are common [7, 8]. Another form of HUS, atypical HUS (aHUS) is distinct from eHUS, while often has overlapping symptoms. aHUS is due to unregulated complement activation, in most cases secondary to identifiable mutant complement regulatory proteins [9]. Its frequency is only about one-tenth of eHUS [10].

Complement activation has been demonstrated in both eHUS and aHUS [11] and anti-C5 therapy with the monoclonal antibody eculizumab, which blocks the terminal pathway of complement activation, has become a mainstay in the management of aHUS. However, consistent benefit of eculizumab in eHUS is lacking [12] despite anecdotal positive reports [13]. Since inhibition of C5 and the terminal pathway of complement is insufficient, we hypothesized that the lectin pathway, which lies upstream from the terminal pathway, is primarily involved, with subsequent downstream activation of the alternative and terminal pathways. Given that sequence of events, we further hypothesized that blockade of the lectin pathway would be effective in treating eHUS. The lectin pathway, one of three known molecular pathways of complement activation (classical, alternative and mannose-binding lectin (MBL)), also plays an important role in innate immune protection against *A. fumigatus* in immunocompromised patients [14], the pathophysiology of atherosclerosis in humans [15], myocardial infarction, coagulation, brain ischemic injury, and the innate immune response to pneumococcal infection in mice [16–18]. We previously demonstrated that injection of (Stx-2 leads to fibrin deposition in mouse glomeruli that was largely blocked by the co-injection of anti-MBL-antibody 3F8 [19, 20]. However, the composition of thrombi in the affected mouse glomeruli was not delineated in those studies. In the present work, we show that injection of Stx-2 in our mouse model leads to the increase in glomeruli not only of fibrin, but also platelets, consistent with the generation of platelet-fibrin thrombi. Importantly, administration of 3F8 with Stx-2 reduces both platelet and fibrin levels to control levels. We also demonstrate the presence of platelets in kidney of humans with eHUS. This is important because we are showing that platelet-fibrin thrombi may underlie the poorly understood pathophysiology of human eHUS.

## Methods

### Mouse Model and Treatment Groups

To investigate the effect of 3F8 on markers of thrombosis and endothelial cells we used sections of kidney harvested from our mouse model that expresses human MBL2 (MBL2 KI) and lacks murine MBLs (MBL-2+/+ Mbl-A/C−/−). Animals were assigned to one of three groups-- a control group that received intraperitoneal phosphate buffered saline (PBS, 200 μl), a Stx-2 group that received 125 pg/g STX in PBS intraperitoneally and a STX-2/3F8 group that received 30 ug/g of anti-MBL2 antibody in PBS intraperitoneally 12 hours before STX-2 injection. Mice were anesthetized with isoflurane and exsanguinated via cardiac puncture on day 4 of the post-injection observation period. All efforts were made to minimize suffering. Kidneys were snap-frozen in Optimal Cutting Temperature (OCT, Sakura Finetek, USA) compound and used for the preparation of frozen sections. There were five different sets of mice receiving one of the three treatments, with seven to ten experiments involving these sets of mice for each of the studies. Experiments on mice were conducted according to the rules of the Brigham and Women’s Hospital Animal Care and Use Committee (IACUC) and performed under the standards and principles set in the Guide for Care and Use of Laboratory Animals [21]. The study was prospectively approved by the BWH IACUC under protocol #1610.

### Antibodies

The following primary IgG antibodies were used: sheep anti-human/mouse fibrinogen (Thermofisher, Waltham, MA); rat anti-mouse CD41 (Biolegend, San Diego, CA); rabbit anti-mouse von-Willebrand factor (vWF) and rabbit anti-mouse podocin (Sigma, St. Louis, MO). All secondary IgG class antibodies were from Thermofisher (Waltham, MA): donkey anti-sheep Alexa 647, donkey anti-rat Alexa 488, goat anti-rabbit Alexa 488, and goat anti-rabbit Alexa 647.

### Kidney Sections

Freshly cut kidney sections (5 μm, two sections per glass slide) were air dried for 15 minutes, washed twice with PBS (pH 7.4), blocked with 5 % goat serum (Genetex, Irvine, CA) for 30 minutes and incubated with primary antibodies for 1 hour at room temperature. In one set of experiments sheep anti-human/mouse fibrinogen (dilution 1:200) and rat anti-mouse CD41 (dilution 1:100) were used. In another set of experiments, the above anti-fibrinogen antibody plus a rabbit anti-mouse vWF (dilution 1:200) were used. In control experiments rabbit anti-mouse podocin (dilution 1:1,000) was used. Primary antibodies were prepared in PBS with 5% goat serum. All sections were washed three times for 3 minutes each with PBS/5% goat serum and then incubated with secondary antibodies in PBS/5% goat serum. For secondary antibodies, we used a donkey anti-sheep Alexa 647 (dilution 1:500) for fibrinogen detection; a donkey anti-rat Alexa 488 (dilution 1:2,500) for CD 41 detection; and, a goat anti-rabbit Alexa 488 (1:5,000) for vWF and podocin detection. Sections were washed with PBS/5% goat serum 3 times for 3 minutes each and fixed with Aqua-Mount (Polysciences, Warrington, PA). Control kidney sections lacked primary antibodies.

### Digital Image Analysis

Kidney sections, covered with cover-slips, were examined using a Nikon Optiphot microscope, with 10x and 20x fluorescent objectives. High-resolution digital images (1,447,680 pixels per image) were taken using a Photometric CoolSnap HQ camera (4095 grey levels, Roper Scientific), a Lumen 200 PRO light source (Prior), and MetaMorph Premier software (Molecular Devices, Inc). Each image at 20x corresponded to an area of a given frozen section of 0.25 mm^2^. Images (two per section, four per slide of two sections) were analyzed for the distribution and co-distribution of the tested markers above, using regions of interest defined as the total area contained within circular approximations of the glomeruli found in a given image. The percent pixels positive for a given marker were averaged for each set of four images. Kidney sections were immunostained concomitantly for each triad of animals receiving control, Stx-2, or Stx-2 plus 3F8 treatment. Data for each animal triad were normalized by the value of pixels positive for each marker for the animal receiving control treatment, giving rise to relative values for pixels positive. Image pixel positive for fibrin(ogen) and CD41 were expressed both as normalized pixel numbers, and as ratios of normalized pixel numbers to number of CD41-positive objects. Ratios of fibrin(ogen) and CD41 pixels to number of CD41-positive objects were taken to be measures of fibrin associated with a given platelet aggregate, and number of platelets in a given platelet aggregate.

### Electron Microscopy

Electron Microscopy (EM) was performed on mouse kidney tissues frozen in OCT and kept at −80°C until sectioned. Freshly processed and frozen untreated C57BL/6 mouse kidneys were used for comparison. Frozen tissues were cut and 1.0 mm sections thawed directly into Karnovsky’s KII Solution (2.5% glutaraldehyde, 2.0% paraformaldehyde, 0.025% calcium chloride, in a 0.1M sodium cacodylate buffer, pH 7.4), fixed overnight at 4°C, and stored in cold buffer. Then the tissue was placed into fresh EM fixative (Karnovsky’s KII Solution) at room temperature for 3 hours. Processing was done in an EMS (Electron Microscopy Sciences) Lynx™ automatic tissue processor. Briefly, tissue was post-fixed in osmium tetroxide, stained *en bloc* with uranyl acetate, dehydrated in graded ethanol solutions, infiltrated with propylene oxide/Epon mixtures, flat embedded in pure Epon, and polymerized overnight at 60°C. One micron-thick sections were cut, stained with toluidine blue, and examined by light microscopy. Representative areas were chosen for electron microscopic study and the Epon blocks were trimmed accordingly. Thin sections were cut with an LKB 8801 ultramicrotome and diamond knife, stained with Sato’s lead, and examined in a FEI Morgagni transmission electron microscope. Images were captured with an AMT (Advanced Microscopy Techniques) 2K digital CCD camera. **Review of Clinical Samples of eHUS**

In our review of clinical samples, we identified 2 cases of eHUS. Archived Epon blocks of the clinical cases were recut, stained and examined for light and electron microscopic studies as described above.

### Statistical Analysis

A two-sided ANOVA was employed for analysis of the image pixels positive for a given antigen following treatment with either Stx-2 or Stx-2 plus 3F8, normalized by the image pixels positive for control (no treatment). Differences were considered significant if the two-sided probability (p), after Bonferroni correction, was less than 0.05, 0.01, or 0.001.

## Results

### Podocin and CD41 in Glomeruli

To identify glomeruli, which are globular structures in frozen sections viewed by IF or phase contrast microscopy, we used anti-podocin antibody, to confirm the presence of podocytes. (Figure 1A). The diameter of glomeruli ranged from 70 to 80 microns, the known diameter of mouse glomeruli [22, 23]. Phase contrast images of the same kidney sections (that were stained for IF) showed structures of similar diameter. CD41, a marker of platelet glycoprotein IIb (GPIIb), was positive within glomeruli (Fig 1B, 1C).

**Fig 1.**
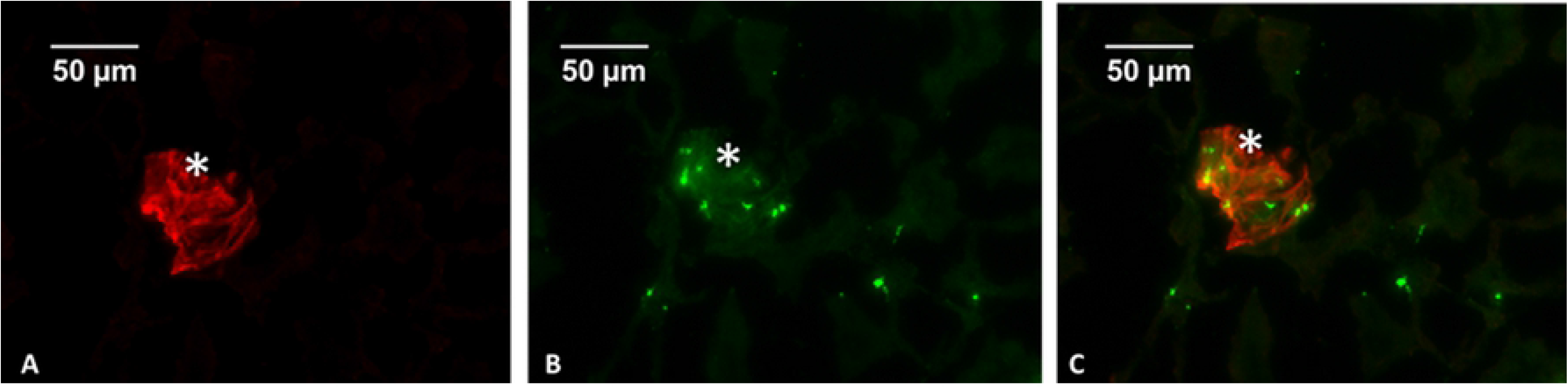
Double immunofluorescence of mouse kidney sections using anti-podocin (A, red) and anti-CD41 (B, green) antibodies, and the overlay (C). The glomerulus (asterisk) is delineated by podocin (magnification 20x, scale bars represent 50 microns).

### Fibrin(ogen) and CD41 in Glomeruli

Both glomerular fibrin(ogen) (p = 0.0023) and CD41 (p = 0.0151) were significantly increased after Stx-2 administration (Figs. 2 and 3). When 3F8 was administered with Stx-2, both fibrin(ogen) (p=0.0061) and CD41 (p = 0.0075) were significantly reduced towards baseline (Fig. 2 and 3).

**Fig 2.**
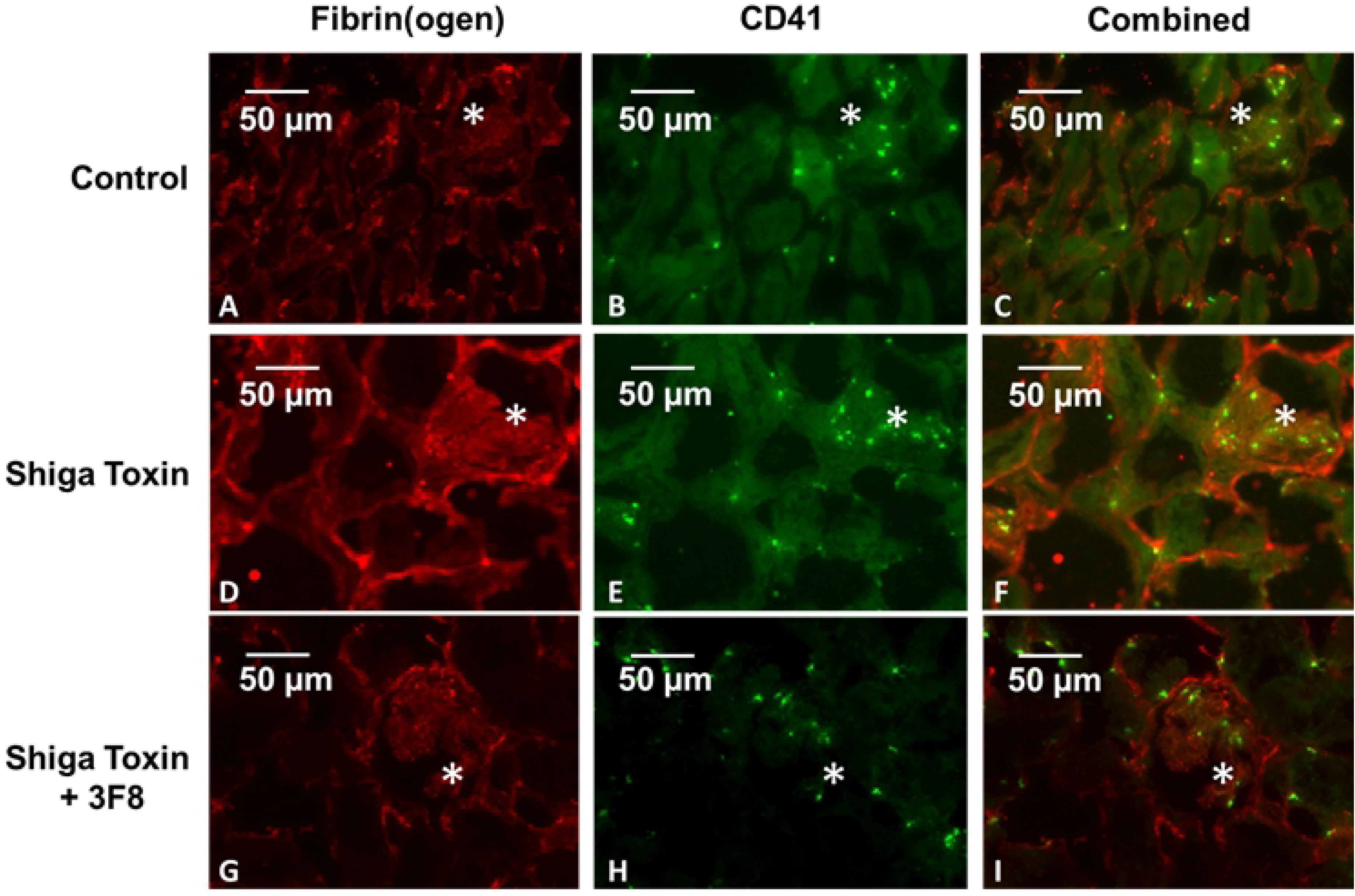
Double immunofluorescence of mouse kidney sections using anti-fibrin(ogen) and anti-CD41 antibodies in control (A-C), Stx-2 (D-F), and Stx-2 + 3F8-treated (G-I) animals. There are significant increases in CD41 (E), fibrin(ogen) (D), and both (F) within the glomerulus (asterisks) after Stx-2 treatment. CD41 (G), fibrin(ogen) (H), and both (I) in glomeruli after treatment with 3F8 return to baseline. There is also some minor CD41-fibrin(ogen) co-localization (yellow) (magnification 20x, scale bars represent 50 microns).

**Fig 3.**
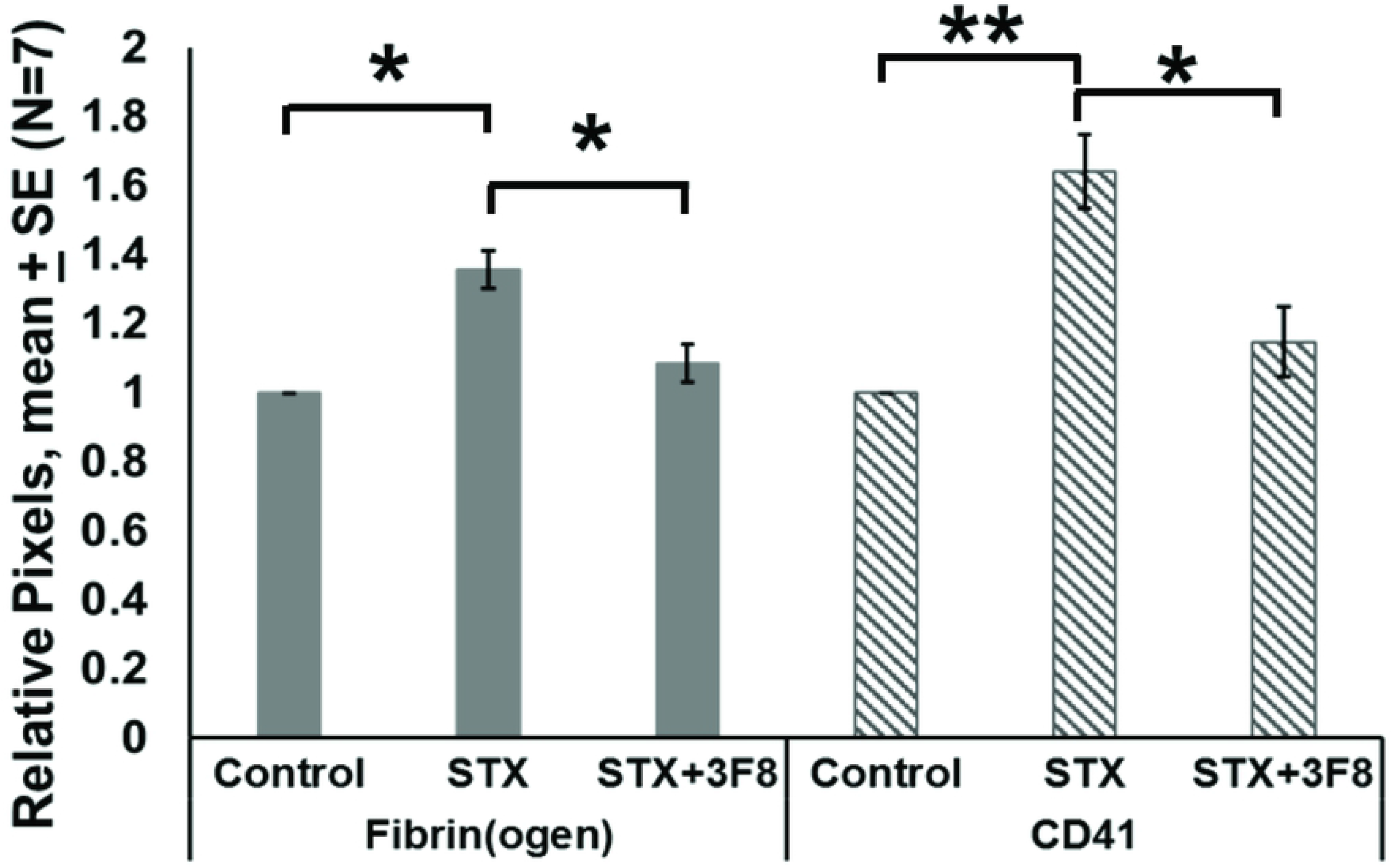
Relative pixels positive for glomerular fibrin(ogen) and CD41 vs. treatment. Mean ± SE, N = 7, *p<0.05, **p<0.01, ***p<0.001.

Thus, 3F8 blocked the Stx-2 induced increases in both fibrin(ogen) and platelets. For control animals, we assume that the CD41 positive objects are single platelets, since the average number of pixels positive for CD41-positive object ranged from 39 to 120, values in the range that we previously reported for a single human platelet [24]. As mouse platelets are somewhat smaller than human platelets, this range can be expected to be a little lower than that for human. Following treatment with Stx-2, we observed greater CD41-positivity, but this increase in signal was not primarily due to an increase in the number of objects. We interpret this to mean that multiple platelets were now present in platelet thrombi. To estimate the number of platelets comprising the platelet thrombi, we then divided the pixel number associated with CD41 positivity by the number of objects observed. This ratio increased (p = 0.0056) with Stx-2 treatment, as compared to control. Fibrin(ogen) associated with a given object, (pixels positive for fibrin(ogen) divided by the CD41-positive object number), also increased (p < 0.0001). Treatment with both 3F8 and Stx −2, however, led to decreases in both platelets per object (p = 0.0112) and fibrin(ogen) per object (p =0.0016), as shown in Fig. 4.

**Fig 4.**
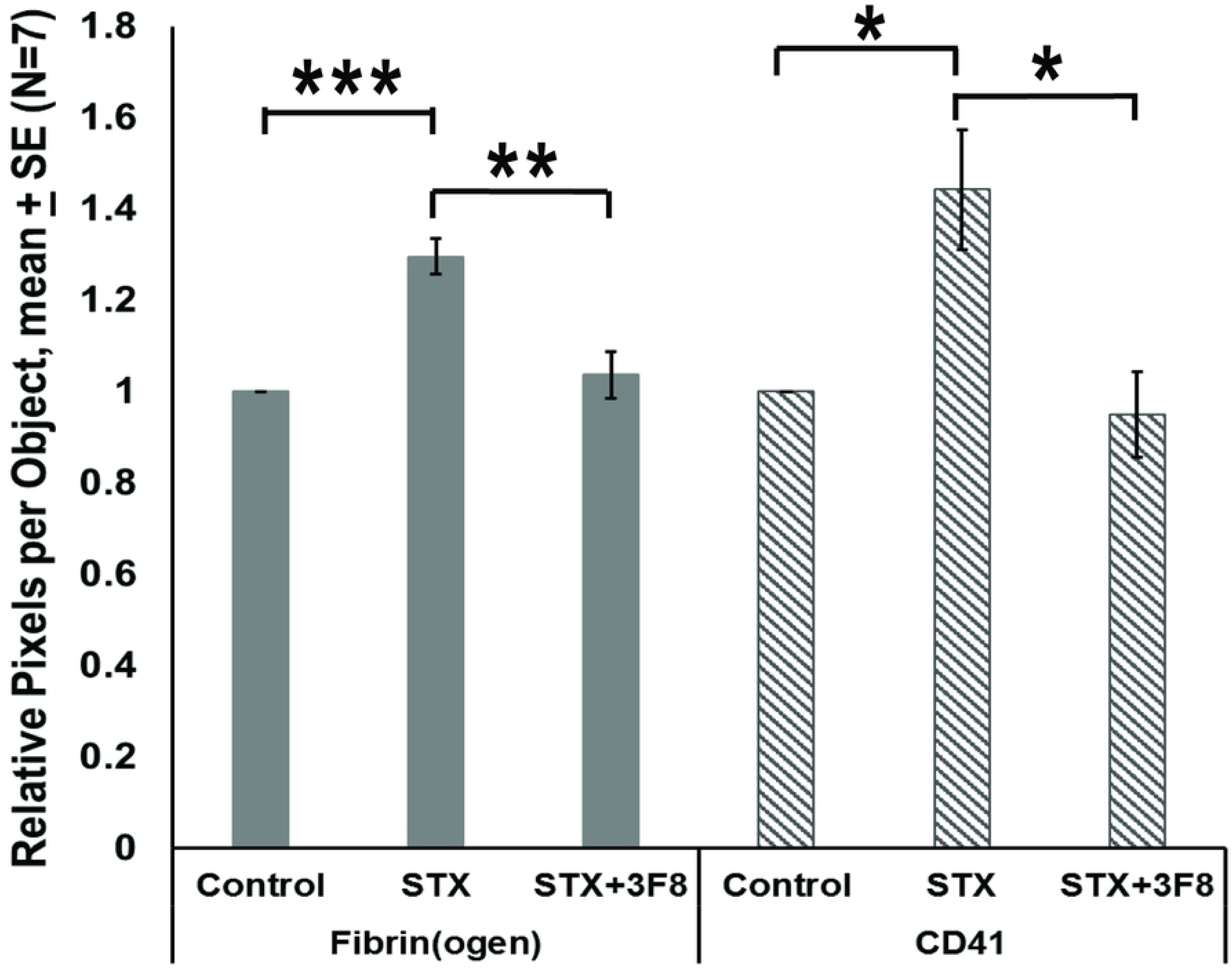
Relative pixels per CD41 positive object for glomerular fibrin(ogen) or CD41 vs. treatment. Mean ± SE, N = 7. *p<0.05, **p<0.01, ***p<0.001.

### Electron Microscopy of Glomeruli

Electron microscopy was used to confirm platelets presence in kidneys’ glomeruli. It demonstrated platelets in glomerular capillaries of Shiga toxin-treated mice, (Fig. 5A) and, for comparison, in untreated C57BL/6 mice (Fig. 5B). This provides confirmation that the CD41-positive objects were platelets.

**Fig 5.**
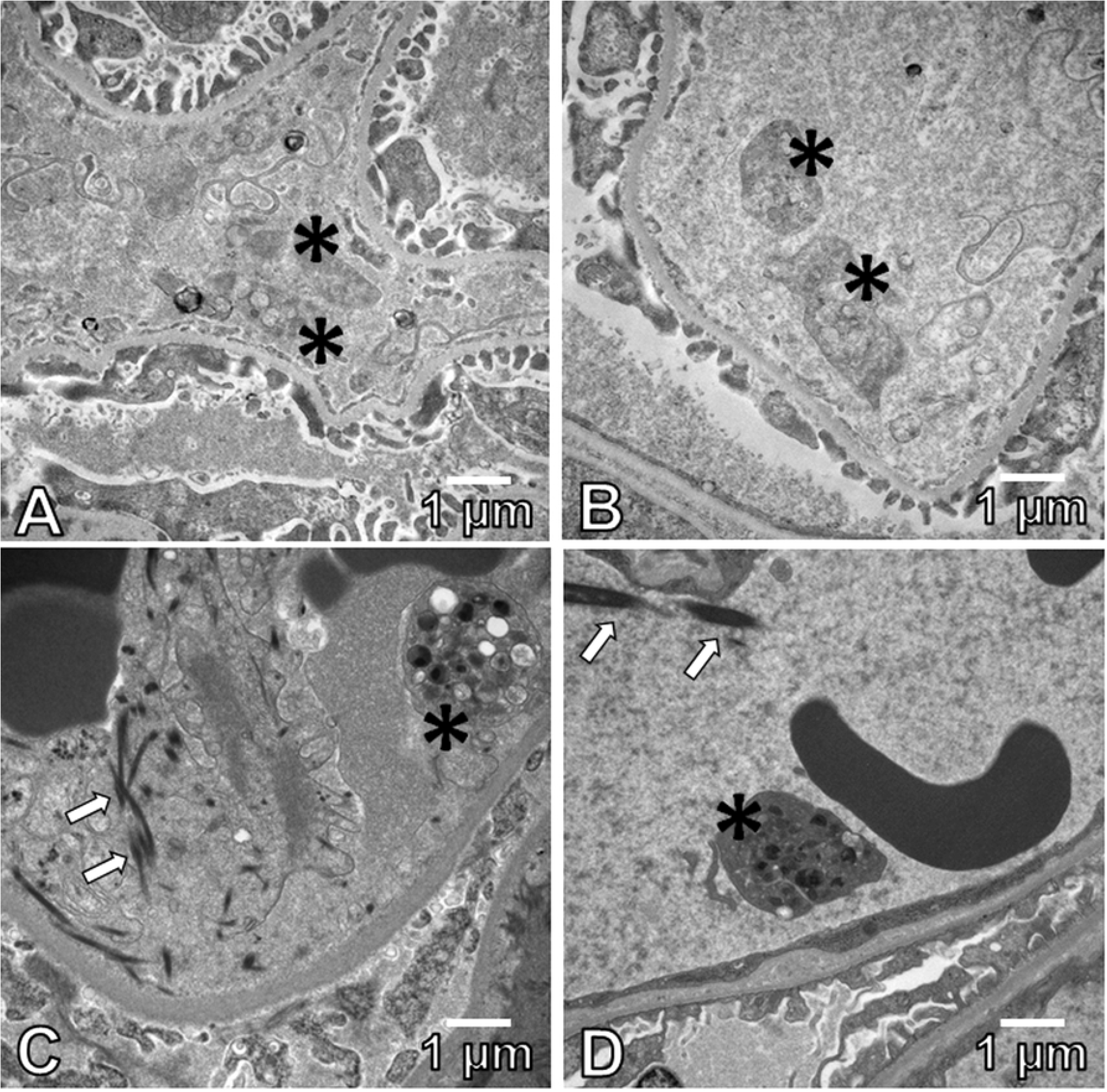
Electron microscopy of platelets (asterisks) in glomerular capillaries of Stx-2-treated mouse (A), untreated C57BL/6 mouse (B) and human kidney biopsies (C, D) in 2 cases of clinical eHUS (11,000x). Arrows denote fibrin tactoids (magnification 11,000, scale bars represent 1 micron).

### Pathology in Clinical eHUS

The presence of platelets and fibrin was also demonstrated in human kidneys. Two cases of pediatric eHUS, in 2 and 3-year-old girls, respectively, presented with gastrointestinal symptoms, severe hemolysis and oligoanuric kidney failure. Kidney biopsies from both showed diffuse acute thrombotic microangiopathy in glomeruli. Electron microscopy in both cases also showed platelets (and fibrin tactoids) in the glomerular capillaries (Fig. 5C, 5D).

### von-Willebrand Factor and Fibrin(ogen) in Glomeruli

Double immunofluorescence showed that vWF, a marker of vascular endothelial cells and platelets, was present in glomeruli (Fig. 6). Glomerular fibrin(ogen) and vWF both significantly increased after Stx-2 treatment (fibrin(ogen) p = 0.0005 and vWF,p = 0.0014). When 3F8 was given with Stx-2, fibrin(ogen) levels were reduced towards baseline (p = 0.0032); however, vWF was not significantly decreased after Bonferroni correction. (Fig. 7).

**Fig 6.**
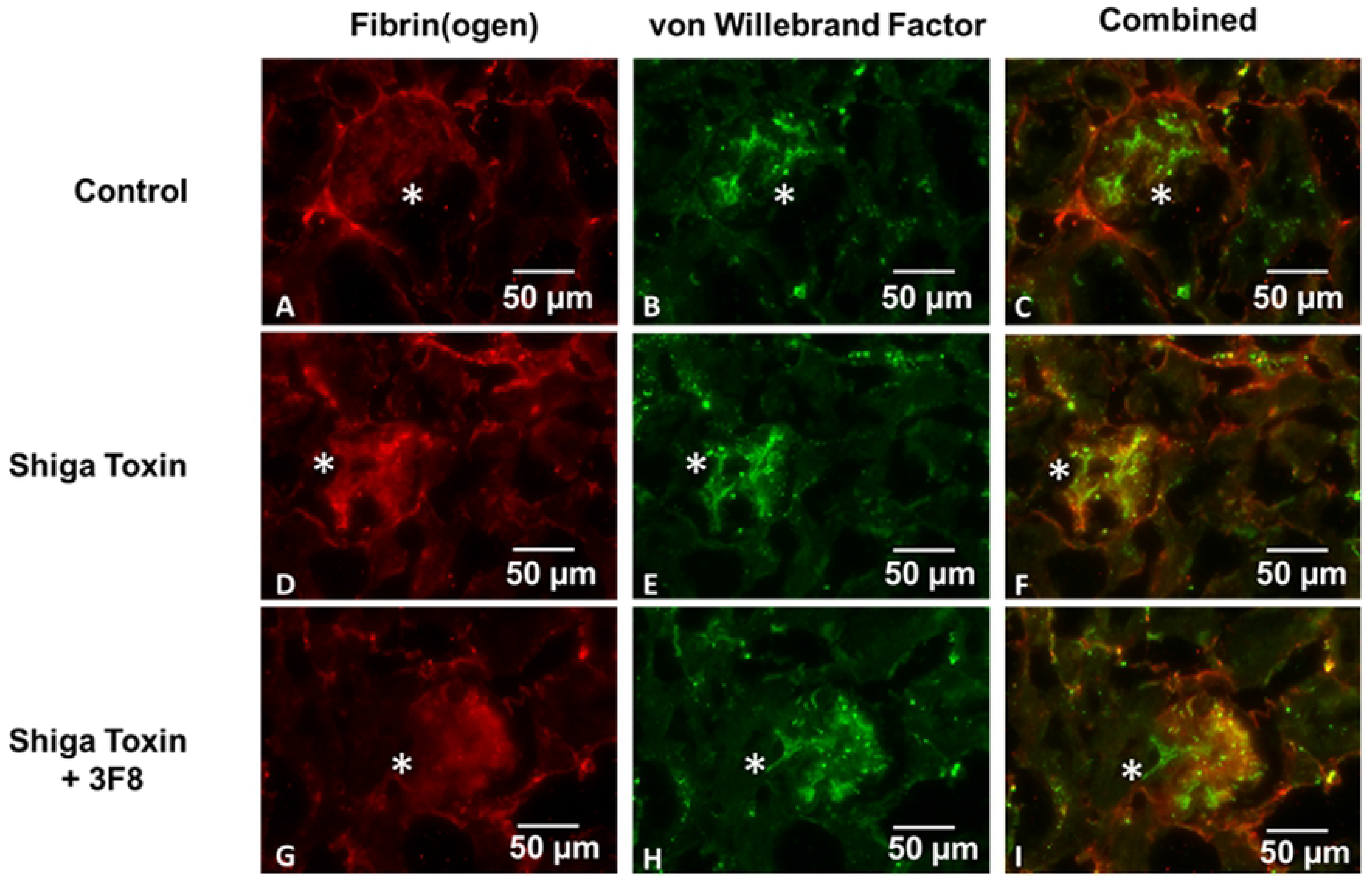
Double immunofluorescence of mouse kidney sections using anti-fibrin(ogen) and anti-vWF antibodies in control (A-C), Stx-2 (D-F), and Stx-2 + 3F8-treated (G-I) animals. vWF-fibrin(ogen) co-localization (yellow) and co-distribution are seen in glomeruli (asterisks). Fibrin(ogen) in glomeruli after treatment with 3F8 (I) returns to baseline (magnification 20x, scale bars represent 50 microns).

**Fig 7.**
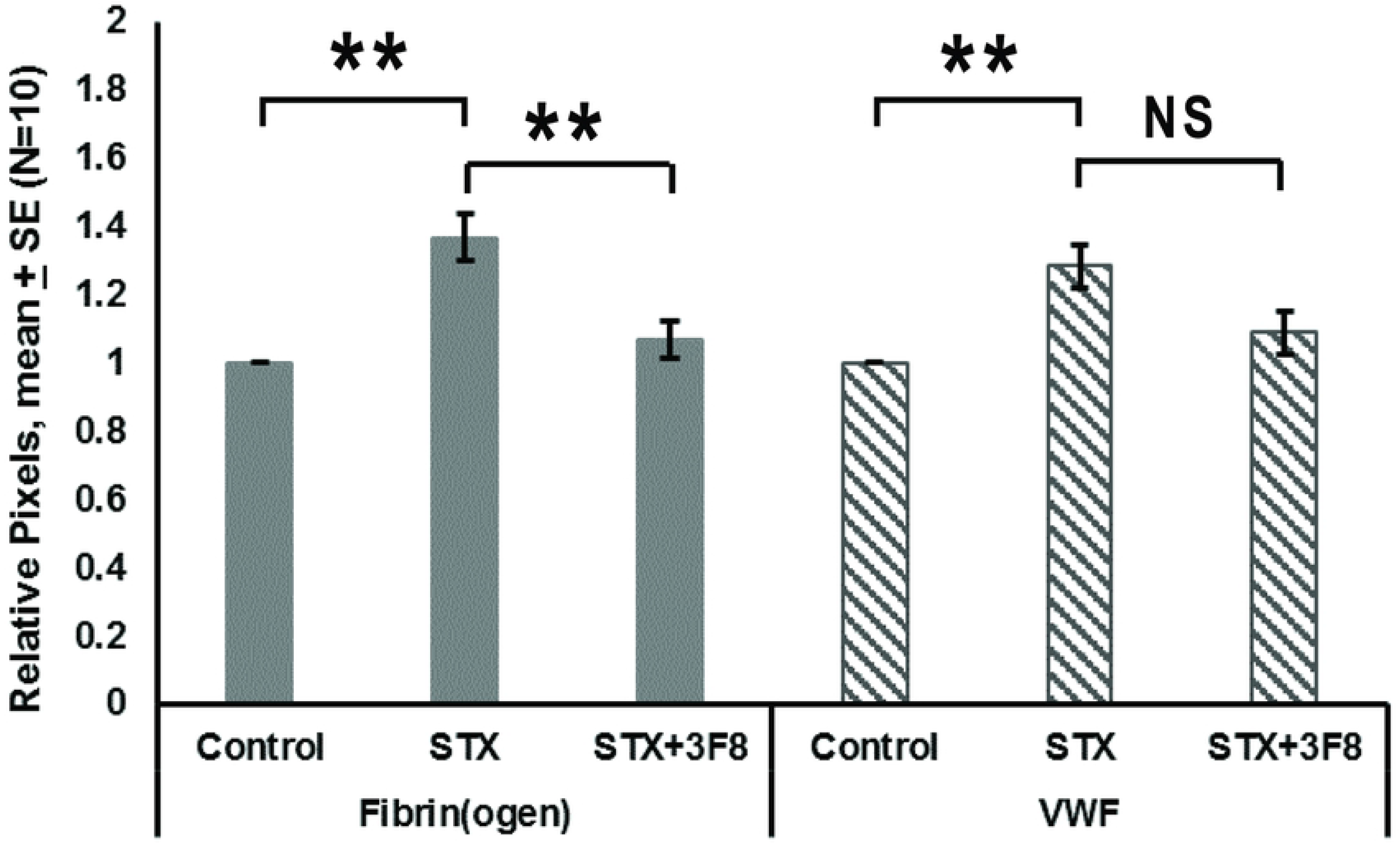
Relative pixels positive for glomerular fibrin(ogen) and vWF vs. treatment. Mean ± SE, N= 10, *p<0.05, **p<0.01, ***p<0.001.

## Discussion

The present study demonstrates that the lectin pathway of complement is important in the development of an eHUS-like state in our murine model of eHUS. In mouse glomeruli, we observed an increase in both platelets and fibrin. With co-administration of 3F8, however, the increases in platelets and fibrin were both prevented, as were clinical and laboratory signs of the eHUS-like state [19], which included impairment of renal function, a decrease in platelet count, and evidence of a thrombotic microangiopathy.

Stx, the triggering agent of eHUS, consists of an enzymatically active A subunit non-covalently associated with five identical B subunits. The B subunit binds to cell membranes that express globotriaosylceramide (Gb_3)_ receptors on human glomerular endothelium and the epithelium of proximal tubules in humans [25, 26] and mice [27], and on epithelium of the gastrointestinal tract and brain microvascular endothelium in humans [28, 29]. The A subunit then enters the cell, where it injures the eukaryotic ribosome, interfering with protein synthesis in target cells. That action, ultimately, alters the expression of certain proteins, including tissue factor and tissue factor pathway inhibitor [30, 31], and induces apoptosis. Stx-producing infection also contributes to release of inflammatory cytokines and cytokine-mediated events, particularly TNFα [32], platelet activation [33], increased procoagulant tissue factor activity on glomerular endothelial cells [34], and activation of complement [19, 20, 35]. In our mouse model, as in all published mouse models, Gb3 is not expressed on glomerular endothelium, although it is expressed on proximal tubules and other tissues, as already noted. Nonetheless, we have determined an important role for the lectin pathway of complement in our model, resulting in systemic activation of complement and consequent development of platelet thrombi in mouse glomeruli, which in turn is associated with kidney injury.

There is growing evidence from in studies on patients with eHUS that one or more complement pathways are activated during the illness [2, 11, 36]. *In vitro* studies demonstrate that Stx-1 and Stx-2 activate the alternative complement pathway leading to increased terminal complement complex C5b-9 [37] and that alternative pathway-deficient mice have reduced renal injury following Stx-1/lipopolysaccharide (LPS) injection [35]. These clinical and *in vitro* investigations, however, did not seek and therefore do not exclude an upstream activation of the lectin pathway. If activated, the lectin pathway would be able to secondarily activate the alternative pathway. The effects of MBL blockade in eHUS are highly relevant, because the lectin pathway can independently enhance clot formation via mannin-binding serine protease-1 (MASP-1) [38].

Importantly, Stx may activate the lectin pathway through the binding of Stx to Gb_3_ with subsequent generation of reactive oxygen species, which are required for MBL ligand expression on human glomerular endothelial cells [39, 40]. Recently we [19, 20] showed that MBL2 plays a crucial role in eHUS. Mice that received intraperitoneal Stx-2 injection developed increases in serum Cr and cystatin C, along with increased free hemoglobin levels, decreased platelet counts, and decreased deposition of C3d in the renal proximal tubule. When 3F8 was administered with Stx-2, plasma MBL2 levels were virtually eliminated and serum Cr and cystatin C levels were preserved at or near control levels, and other effects of Stx-2 were prevented. Thus, inhibition of the lectin pathway of complement significantly protected the mice against the complement-mediated renal injury induced by Stx-2.

In our mouse model Stx-2 injection led to the presence of increased platelets and fibrin(ogen) deposition in murine glomeruli, together with an increase in the size of the CD41-positive objects and therefore, we infer, an increase in the number of platelets. Thus, the platelet-fibrin thrombi in our model are characterized by an increased number of platelets, consistent with the presence of platelet aggregate formation. The aggregates, in turn, have associated fibrin(ogen) in proportion to the number of incorporated platelets. The aggregates, in turn, have associated fibrin(ogen) in proportion to the number of incorporated platelets, but, we speculate, not increased to a level that might suggest the local secretion from endothelium of high MW vWF multimers. These increases were prevented by 3F8. We interpret our findings to be consistent with the increased and then blocked formation of glomerular platelet-fibrin thrombi. In the present study, we used anti-fibrin(ogen) antibody for fibrin detection, in contrast to our previous work (19) in which the presence of fibrin in glomeruli was detected using an anti-fibrin antibody and an immunoperoxidase technique. To our knowledge, only the present anti-fibrin(ogen) antibody is available for double immunofluorescence analysis. In the prior study (19), little or no fibrin was detected outside of glomeruli, but in our current work, fibrin(ogen) signal was present outside of glomeruli, in the tubules and interstitium. The fibrin(ogen) signal outside the glomeruli, we believe, is largely fibrinogen and not fibrin.

Despite a report that Stx induces vWF secretion by human endothelial cells [41], the presence of increased vWF in the glomeruli thrombi of human eHUS is controversial [9, 42]. Kidney tissue from some patients with HUS are reported to have vWF/platelet-rich thrombi [9] but in other studies glomerular thrombi contain fibrin and little or no vWF [42]. In the present work, we found no evidence for increased vWF following Stx-2 treatment in proportion to the number of CD-41-positive objects (platelets). The failure of the vWF to return to baseline after Stx-2 + 3F8, however, could be consistent with Stx-2-induced release of endothelial cell and/or platelet α granule vWF.

The presence of platelets in glomerular thrombi in human eHUS and in animal models of eHUS is unclear. Hosler et al. [43] reviewed 56 autopsy cases of thrombotic thrombocytopenic purpura (TTP, another thrombotic microangiopathy) and HUS and observed platelet-rich thrombi in the kidneys of patients with classical TTP (25 cases) but not in patients with HUS (31 cases), for which only red cell-fibrin thrombi were present. Others [11, 42] have made similar observations. Of importance, these studies did not describe presence or absence of platelets, and none reported EM studies. Two recent papers using mouse models reached conflicting conclusions. Therefore, our work, to our knowledge, is the first to show that the pathophysiology of eHUS may be one of platelet-fibrin thrombi in the glomeruli, as opposed to high MW vWF multimers, as in TTP, or fibrin alone.

Porubsky et al. [27] used transmission EM and reported the presence of thrombotic material composed of platelets and fibrin in glomerular capillaries of patients with Stx-2-associated kidney failure, but not in their mouse model of eHUS. On the other hand, Morigi et al. [35] observed platelets on glomerular endothelium in a murine model of eHUS, although their mice were treated with LPS in addition to Stx-2. Our study demonstrates the presence of platelets in a murine model of eHUS. This is similar to our observations in 2 clinical cases and in the studies by Porubsky et al. [27].

In summary, the present study demonstrates that Stx-2 leads to the formation of glomerular platelet-fibrin thrombi in our mouse model of eHUS. The co-administration of the MBL2 inhibitor 3F8 with Stx-2 prevented platelet-fibrin deposition. These findings suggest a role for the lectin pathway of complement in human eHUS, and, importantly, may provide a promising target for future therapy for this condition.

## Acknowledgements

We are thankful to M. K. Selig for his help with electron microscopy.

